# *Naa15* knockdown enhances c2c12 myoblast fusion and induces defects in zebrafish myotome morphogenesis

**DOI:** 10.1101/306563

**Authors:** Olivier Monestier, Aurélie Landemaine, Jérôme Bugeon, Pierre-Yves Rescan, Jean-Charles Gabillard

## Abstract

The comprehension of muscle tissue formation and regeneration is essential to develop therapeutic approaches against muscle diseases or loss in muscle mass and strength during ageing or cancer. One of the critical steps in muscle formation is the fusion of muscle cells to form or regenerate muscle fibres. To identify new genes controlling myoblast fusion, we undertook an siRNA screen in c2c12 myoblasts and found that N-alpha-acetyltransferase 15 (*Naa15*) knockdown enhanced c2c12 myoblast fusion suggesting that *Naa15* negatively regulated myogenic cell fusion. We identified two *Naa15* orthologous genes in zebrafish genome: *naa15a* and *naa15b*. These two orthologs are both expressed in myogenic domain of the somite. Knockdown of zebrafish *naa15a* and *naa15b* genes induced a “U” shaped segmentation of the myotome and alteration of myotome boundaries resulting in the formation of abnormally long myofibres spanning adjacent somites. Taken together these results show that Naa15 regulates myotome formation and myogenesis in fish.

## 1. Introduction

The fusion process is a critical step for the formation and reparation of muscle. Indeed, skeletal muscle is mainly composed of multinucleated cells, called myofibres. To form those myofibres, progenitor cells (myoblast) proliferate, differentiate into myocytes, fuse to form multinucleated myotubes and finally mature into functional myofibres. The process leading to the formation of skeletal muscle is usually divided into a primary, a secondary, and an adult phases (1). The primary phase is powered by cells originating from dermomyotome lips and results in the formation of the primary myotome. Then, during the secondary phase, cells emanating from the central region of the dermomyotome, differentiate and fuse either with each other to form secondary fibres (hyperplasic growth) or with primary fibres (hypertrophic growth). Through adult phase, satellite cells allow muscle growth, and regeneration after damage and injury.

Myocyte fusion is a very coordinated event requiring that two cells get close, recognize each other, adhere their membrane, open fusion pore and finally merge together into one multinucleated cell (2). This process implies many known molecular components studied in multiple model organisms like drosophila, zebrafish and mouse.

In the fruit fly *Drosophila melanogaster*, each muscle is composed of a single myofibre formed by the fusion of a unique founder cell (FC) with fusion competent myocyte (FCM). Recognition and adhesion between FC and FCM are mediated by immunoglobulin domain containing cell adhesion molecules. Among them, are Kin of IrreC (Kirre) and Roughest (Rst) which are expressed in FC (3,4) as well as ticks and stones (Sns) and Hibris (Hbs) both expressed in FCM (5–8). In vertebrate species the presence of two types of muscle cells has not yet been demonstrated, nevertheless, some genes initially identified in drosophila have homologs in vertebrates. For example Kirrel (Kirre homologue) and nephrin (Sns homologue) are both involved in cell recognition/adhesion process (9,10). The fusion of myocytes in vertebrates also implies specific factors such as myomaker, myomerger and myomixer (11–15) and some, species-specific factors such as Jamb and Jamc in zebrafish (16) or Itgb1 in mouse (17).

After the initial recognition/adhesion step, greater membrane proximity is required and reached by a reorganization of actin cytoskeleton. This is achieved, in Drosophila, by regulators like WASP and Scar that affect actin polymerization mediated by the Arp2/3 complex (18–22). In vertebrate, this step is dependent of Dock1 and Dock5 (23,24), Rac1 (23,25), and N-WASP (26).

At last the lipid bilayer needs to be destabilized to allow the cells to merge. Some studies performed in c2c12 cells and in chicken show that the family of brain angiogenesis inhibitor molecules (BAI) play a major role during this step of the myocyte fusion (27–29).

The regulation of myocytes fusion is highly complex and is far to be completely understood. To identify new genes implicated in the myocyte fusion process in vertebrates, we performed an *in vitro* functional screen in c2c12 cell line based on siRNA knockdown. We found that *Naa15* knockdown led to the formation of myotube larger than those found in c2c12 control cells. In line with this observation, knockdown of the two zebrafish orthologous genes *naa15a* and *naa15b* induced the production of giant myofibres spanning two somites in zebrafish embryos suggesting that *Naa15* negatively regulated myofibre formation.

## 2. Materials and Methods

### Zebrafish husbandry

Wild-type zebrafish (Danio rerio) were raised in our facilities (INRA LPGP, Rennes) and maintained under oxygen saturation in a recirculating water system at 27 ± 1 °C, pH 7.5. Zebrafish were exposed to a photoperiod of 16 h light/8 h dark. Fish used in this study were reared and handled in strict accordance with French and European policies and guidelines of the Institutional Animal Care and Use Committee (DDSV approval #35-47).

### siRNA screen

c2c12-MCK:GFP mouse myoblast cells (ATCC-CRL-1772 modified) were maintained at 37°C and seed in 96 wells plates at a density of 10.000 cells/well in Growth Medium (Dulbecco’s modified Eagle’s medium (DMEM), supplemented with 10% heat-inactivated fetal bovine serum (FBS) and Antibiotic-Antimycotic solution (BIOWEST)). Two hundred fifty five genes expressed in proliferating or differentiating C2C12 cells (Moran et al 2002; Tomczack et al., 2003) but with unknown function, were tested in this screen. Each gene was knocking down with two different siRNA (Flexiplate siRNA, Quiagen). Each plate included two negative controls (no siRNA) and two positive controls treated with anti IL4 siRNA. The cells were transfected at day 1 and day 4 with DMEM including 5nM of siRNA and 1μl per well of INTERFERin (Polyplus) and placed in differentiation medium (DMEM containing 2% FBS and Antibiotic-Antimycotic solution). The medium was changed every day. After six days of differentiation, cells were washed two times with phosphate-buffered saline (PBS) before fixing in PBS with 4% paraformaldehyde for 15 min. Then, the cells were permeabilized 3 minutes in PBS with 0.1% TritonX100, stained with 0.1μg/ml of DAPI for 5 minutes and stored at 4°C in the dark.

Image acquisition was made using an HCS Arrayscan VTI (Cellomics/Thermofisher Scientific) with a ZEISS EC plan NEOFluar 10x ON objective and an ORCA-ER 1.00 camera. A macrocommand was edited for visilog 6.7 to monitor automatically the fusion index, mean nuclei number per GFP+ cells, and the GFP+ cells area.

### Cell culture for QPCR analysis

The c2c12-MCK:GFP mouse myoblast cells were maintained at 37°C and seeded in 12 wells plates at a density of 100.000 cells/well in growth medium. *Naa15* was knocking down with two different siRNA (Flexiplate siRNA, Quiagen). The cells were transfected at day 1 with DMEM including 5nM of siRNA and 3.5μl per well of INTERFERin (Polyplus) and place in differentiation medium. Each plate includes 4 wells transfected with anti-GFP siRNA as negative control, and 4 wells transfected with each of the 2 anti-*Naa15* siRNA. The medium was changed every day. One plate was stopped after 0, 1, 2 or 3 days of differentiation. Total RNA was isolated using TRIzol (Invitrogen) according to the manufacturer’s instructions and relative RNA concentration was determined by spectrophotometric analysis. 0.3 μg of total RNA was reverse-transcribed into cDNA using the high-capacity cDNA reverse transcription kit (Applied Biosystems). Realtime PCR was performed in duplicate using 1/10, 1/40 or 1/400 dilution of RT-cDNA from c2c12 cells. Samples were amplified in a 96 well plate using SYBR Green on a StepOnePlus Real Time PCR system (Applied Biosystems) with specific primers for mouse *Naa15*, and *Myog* genes at a concentration of 300nM. The expression of *Hprt* and B-Actin genes were used as endogenous control to normalize each sample. Relative mRNA expression was assessed by relative standard curve method.

### In situ hybridization

Embryos were removed from their chorion by a 3 min incubation in a 1/100 (wt/vol) Pronase solution (Sigma, P6911) pre-warmed to 28 °C. Then, they were fixed in 4% paraformaldehyde overnight and stored in methanol at −20 °C. Anti-sense RNA probes labelled with digoxigenin (DIG) were prepared from PCR-amplified templates using appropriate RNA polymerases. Whole mount in situ hybridization were performed with an INSITU PRO VS automate (INTAVIS AG) using standard protocol ***(30)***. Whole mount in situ images were obtained using a macroscope NIKON AZ 100 coupled with NIKON Digital Sight DSRi1 camera and using NIS-Elements D 3.2 software.

### Morpholino injection

Freshly fertilized eggs were injected with morpholinos at one to two-cell stage. Morpholinos (Gene Tools) were dissolved in sterile water at a concentration of 2.5 mM. Anti-*naa15a*, anti-n*aa15b*, anti-p53 and anti-*naa15a* mismatch control morpholinos were used. Anti-Naa15a and anti Naa15b were designed to bind the area of the predicted start codon. Morpholinos sequences were as follows: anti-*naa15a*: TCTTGAGGGTTGTCCACCGCGACTT; anti-*naa15b*: CGGCATCCTGTTCACTCTCTATTTC; anti-*naa15a* mismatch control: TATTGA<u>C</u>GGTTGT<u>A</u>CACC<u>C</u>CGA<u>A</u>TT. 300 to 450 eggs were injected for each experiment. Embryos were injected with approximately 4 nl of morpholinos (8.8 ng) diluted in sterile water with 0.1% phenol red. About the same number of eggs was injected with the anti-naa15a mismatch control. To ensure that phenotype specificity is due to knock down of *Naa15* orthologues and not to nonspecific induction of apoptosis, embryos were co-injected with anti-*p53* morpholino (6.5ng of anti-*naa15a*+6.5ng of anti-*p53*). The injected eggs were cultured at 28 °C, and embryos were fixed in 4% paraformaldehyde overnight at 30h of development. Zebrafish embryos were permeabilized in 0.3% Triton in PBS solution for 3 h. Embryos were stained with a solution including 5μg/ml of WGA-Alexa 488 and TOPRO 3 (1:1000 dilution) overnight. Confocal microscopy images were collected using an OLYMPUS BX61WI FV 1000 microscope and FluoView 3.0 software. Entire images were adjusted for contrast, brightness, and dynamic range using ImageJ software.

### Statistical analyses

For the siRNA screen, statistical analysis was made using a One-way ANOVA (ANalysis Of Variance) with post-hoc Tukey test in SigmaStat 3.5. For QPCR experiment statistical analysis was made using a One-way ANOVA in Past 3.15.

## 3. Results

### Naa15 knockdown enhanced c2c12 myoblast fusion

An *in vitro* functional screen was performed based on siRNA knockdown (KD) in MCK:GFP C2C12 myoblasts. Those myoblasts express GFP under the muscle creatine kinase (MCK) promoter, GFP is therefore expressed only in differentiated myoblasts allowing an easy monitoring of cells differentiation. Our functional screening allowed the identification of genes which *in vitro* knock-down significantly impacted myoblast fusion without affecting the differentiation capacity of the cells, various functional parameters were assessed among which the fusion index, mean nuclei number per GFP+ cells, and the GFP+ cells area.

One of the more strong phenotype was observed after the KD of the N (Alpha)-Acetyltransferase 15 (*Naa15* also referred to as *Tbdn, Narg1, mNat1, NATH*) gene (Fig 1A). The c2c12 cells transfected with two different anti-*Naa15* siRNA exhibited a significant increase in the fusion index. This index (number of nuclei in myotubes / total number of nuclei) was around 75% greater in GFP+ (differentiated) cells after treatment with both *Naa15* siRNA than in control cells (Fig 1B). Specifically, the mean number of cells including more than 10 nuclei after 7 days of differentiation was five times higher for cell cultures treated with anti-*Naa15* siRNA when compared to control cell cultures. No significant difference was observed in the number of small myotubes (<4 nuclei) (Fig 1C).

**Fig1.**
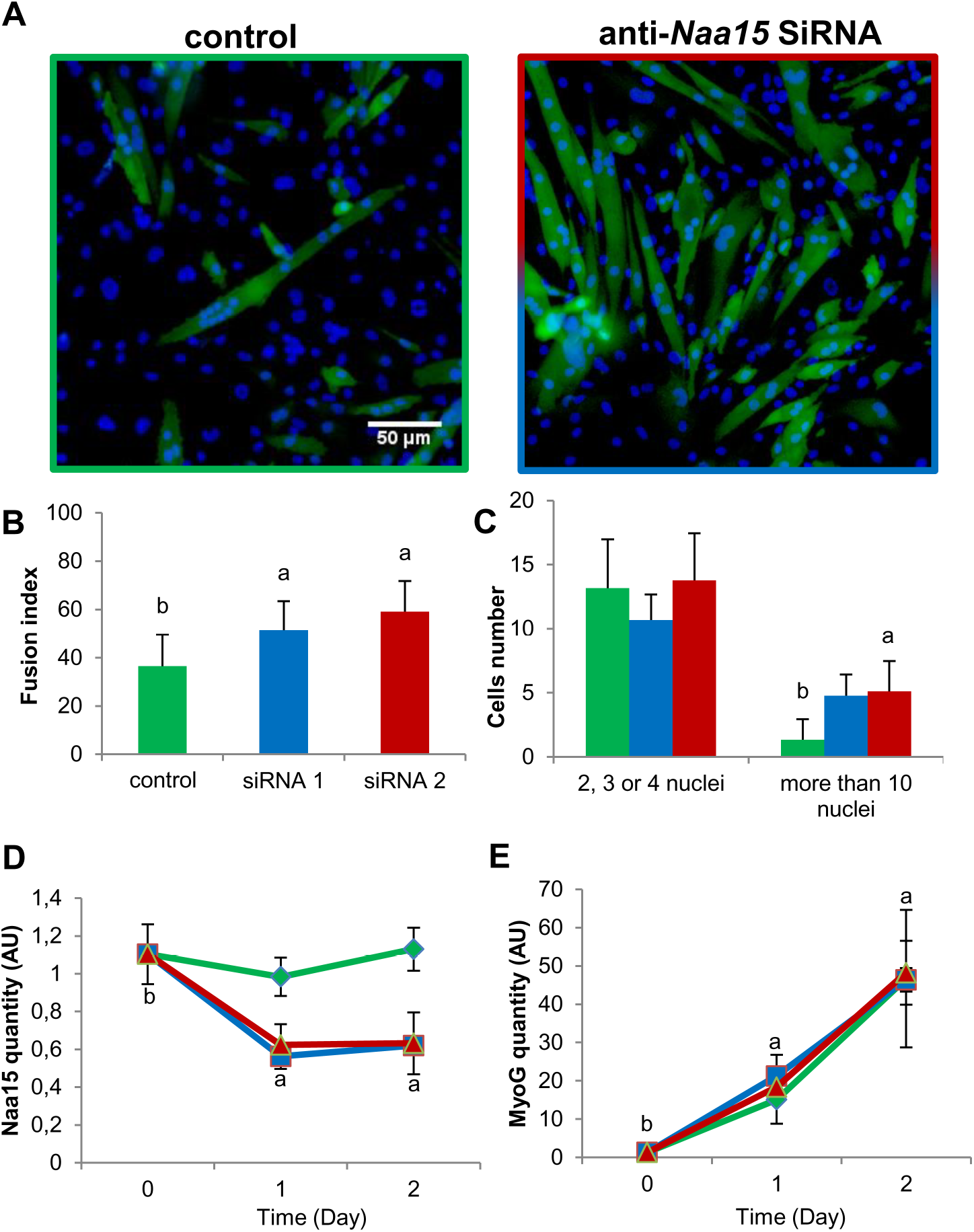
SiRNA screen. *Naa15* act as a fusion inhibitor in c2c12 myoblast. c2c12-MCK:GFP Cell cultures transfected with anti-*Naa15* siRNA and stained with DAPI (blue) present more nuclei in differentiated (GFP+) cells (A). 1B. The fusion index of the GFP+ cells is significantly higher in cultures treated with the anti-*Naa15* siRNA compared to control cell cultures. 1C. The mean number of cells including more than 10 nuclei after 7 days of differentiation is five times higher in cell cultures treated with anti-*Naa15* siRNA compared to control cell cultures. No significant difference is observed in the number of small myotubes (<4 nuclei). QPCR experiment show that, siRNA treatment reduces to half the *Naa15* expression (D). The expression of *Myogenin* (*MyoG*) is not affected by the anti-naa15 siRNA transfection (E). Different letters indicate significant differences between groups (t-test p<0.05). Values represent means ± SD (N=4). Blue and red curves and histograms represent results of cell cultures treated by 2 different siRNA, control group is represented by green curves and histograms.

QPCR experiment showed that both siRNA treatment induced a 50% reduction of *Naa15* expression (Fig 1D). As shown by differentiation index, no significant changes in the expression of *Myogenin* (*MyoG*), a differentiation marker, was observed in the cells transfected with anti-*Naa15* siRNA as compare to controls (Fig 1E). This confirm that the differentiation process was not impacted by *Naa15* knockdown.

### The two Naa15 orthologous genes were expressed in zebrafish somites

To study the function of *naa15* during myogenesis in zebrafish, we looked for orthologous genes. We performed BLAST in public databases and identified two orthologous genes namely *naa15a* and *naa15b*. Those genes encode proteins with respectively 85% and 90% of homology with the murine NAA15 protein. Protein sequences were used to deduce a phylogenetic tree from maximum likelihood method (Fig 2) and show an *naa15* duplication occurring in teleost and in cyprinid as expected.

**Fig2.**
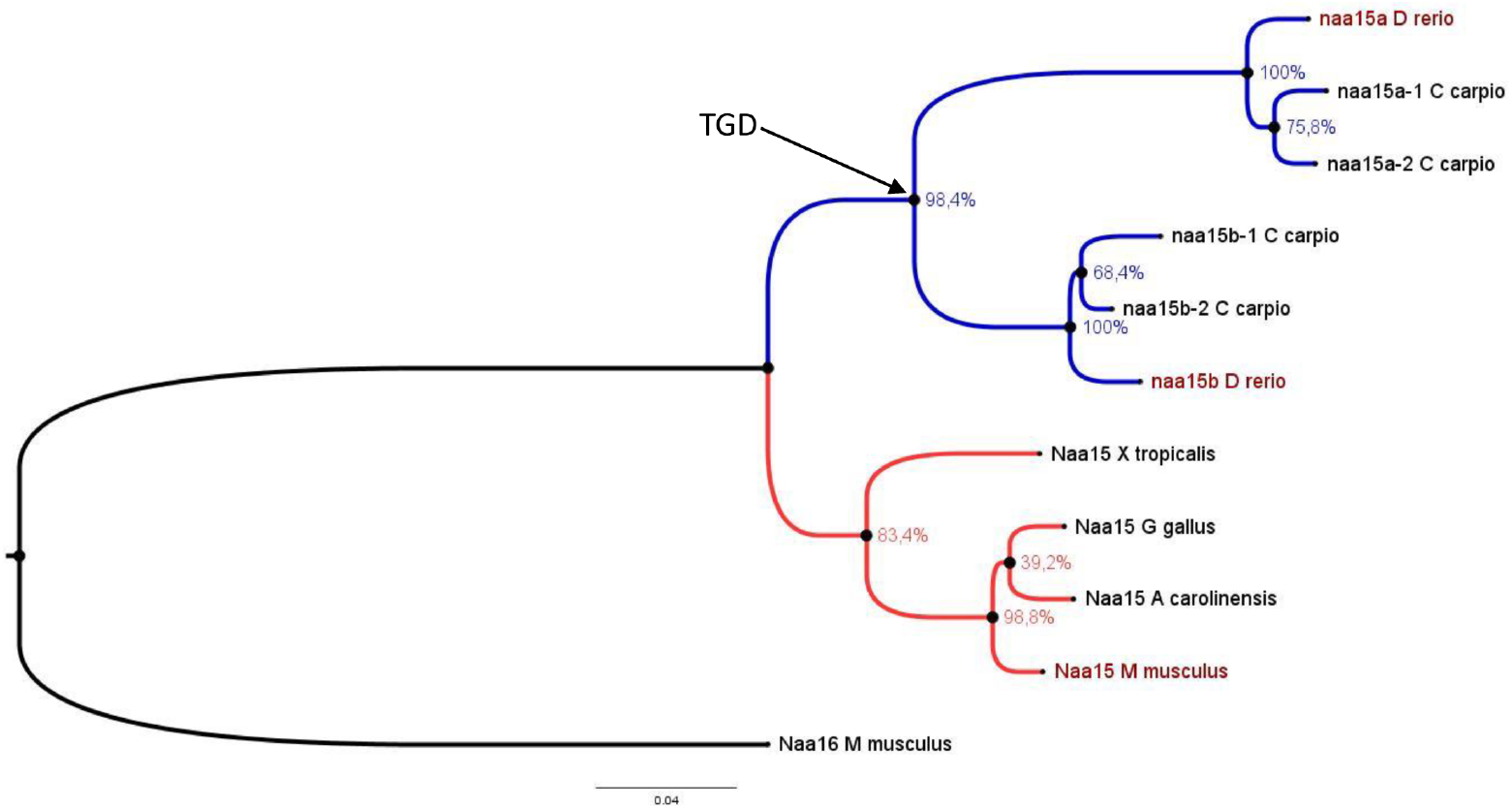
*Naa15* orthologous genes in zebrafish. The evolutionary history of NAA15 related proteins was inferred by using the maximum likelihood method based on the JTT matrix-based model. The bootstrap value calculated out of 500 replicates is indicated for each node. A discrete gamma distribution was used to model evolutionary rate differences among sites. *Naa15* duplicate into *naa15a* and *naa15b* during the Teleost Genome Duplication (TGD).

*In situ* hybridization showed that *naa15a* (Fig 3A) and *naa15b* (Fig 3B) were both expressed in somites during late somitogenesis (24 hpf), when myoblast fusion process occurs. There are also expressed in eyes and midbrain. It could be noticed that only a weak signal was observed for naa15b *in situ* hybridization.

**Fig3.**
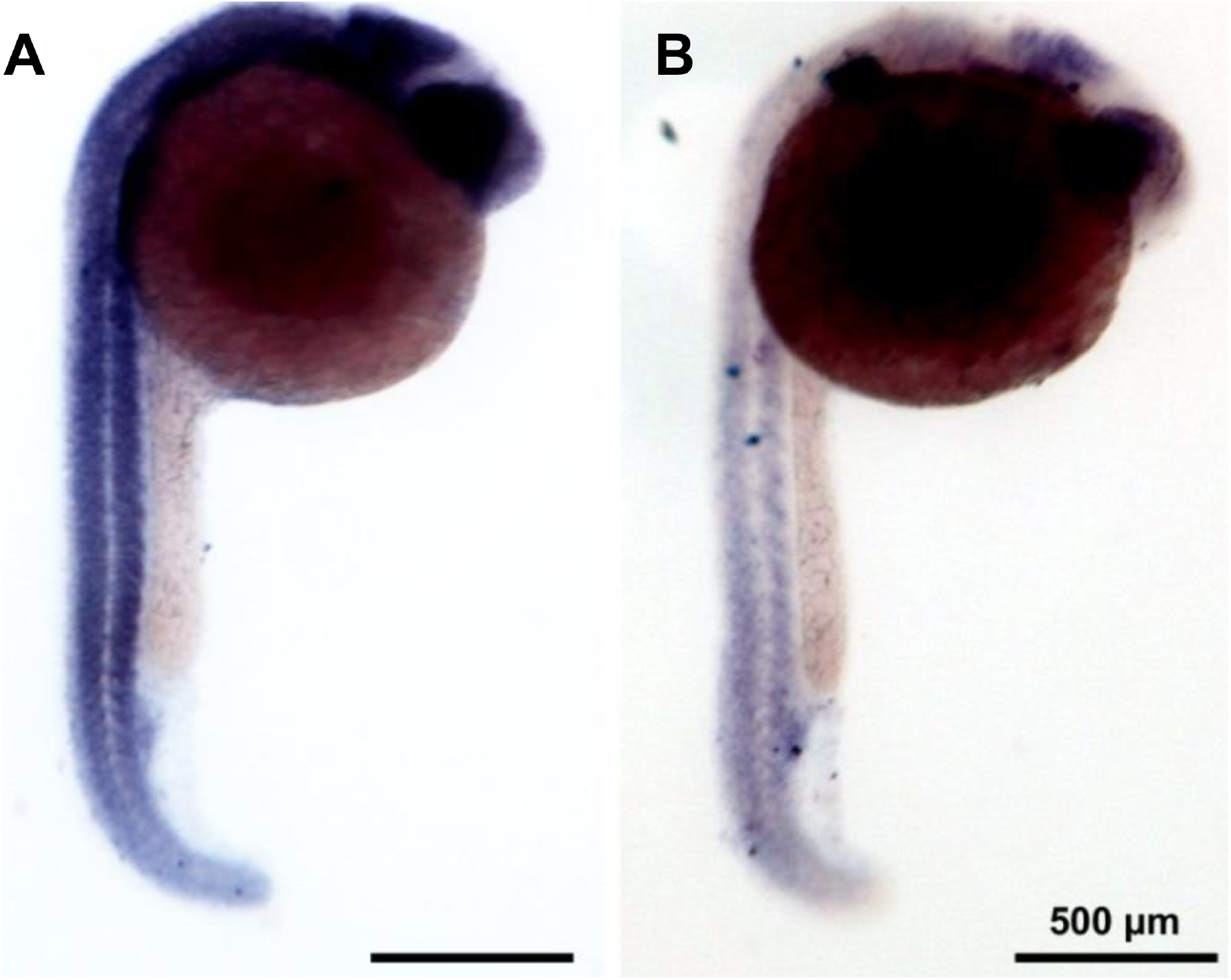
*Naa15a* and *naa15b* are expressed in somite at 24hpf. *naa15a* and *naa15b* expression in 24 hpf zebrafish embryos. *naa15a* is strongly expressed in somites as well as in eyes and midbrain (left panel). *naa15b* is also expressed in somite, eyes and midbrain but the signal appears weaker (right panel).

### naa15a and naa15b knockdown in zebrafish led to curvature of the body

To determine whether *naa15a* and *naa15b* are required for fish myogenesis *in vivo*, we performed morpholino knockdown of *naa15a* and *naa15b*. Zebrafish embryos at 1 cell stage were injected with either an anti-*naa15a*, an anti-*naa15b*, or an anti-*naa15a* mismatch morpholino (4 mismatches) used as a control. The control embryos injected with anti-naa15a mismatch morpholino exhibited a normal morphology at 30hpf and no phenotype was observed as compare to wild type uninjected embryos. At 30hpf, *naa15a* or *naa15b* morphant exhibited morphological defect that could be classified in two classes. The class 1 embryos (C1) had reduction of the body size and small curvature of the trunk. The phenotype of the class 2 embryos (C2) is more severe, and a greater curvature of the trunk is observed (Fig 4). According to the experiment, the C1 embryos represent 30-50% of the *naa15a* and 15-25% of the *naa15b* morphants, and the C2 embryos represent 30-50% of the naa15a and 40-56% of the naa15b morphants. Wild type phenotype is observed in 5-20% of the *naa15a* morphants and in 20-30% of the *naa15b* morphants.

**Fig4.**
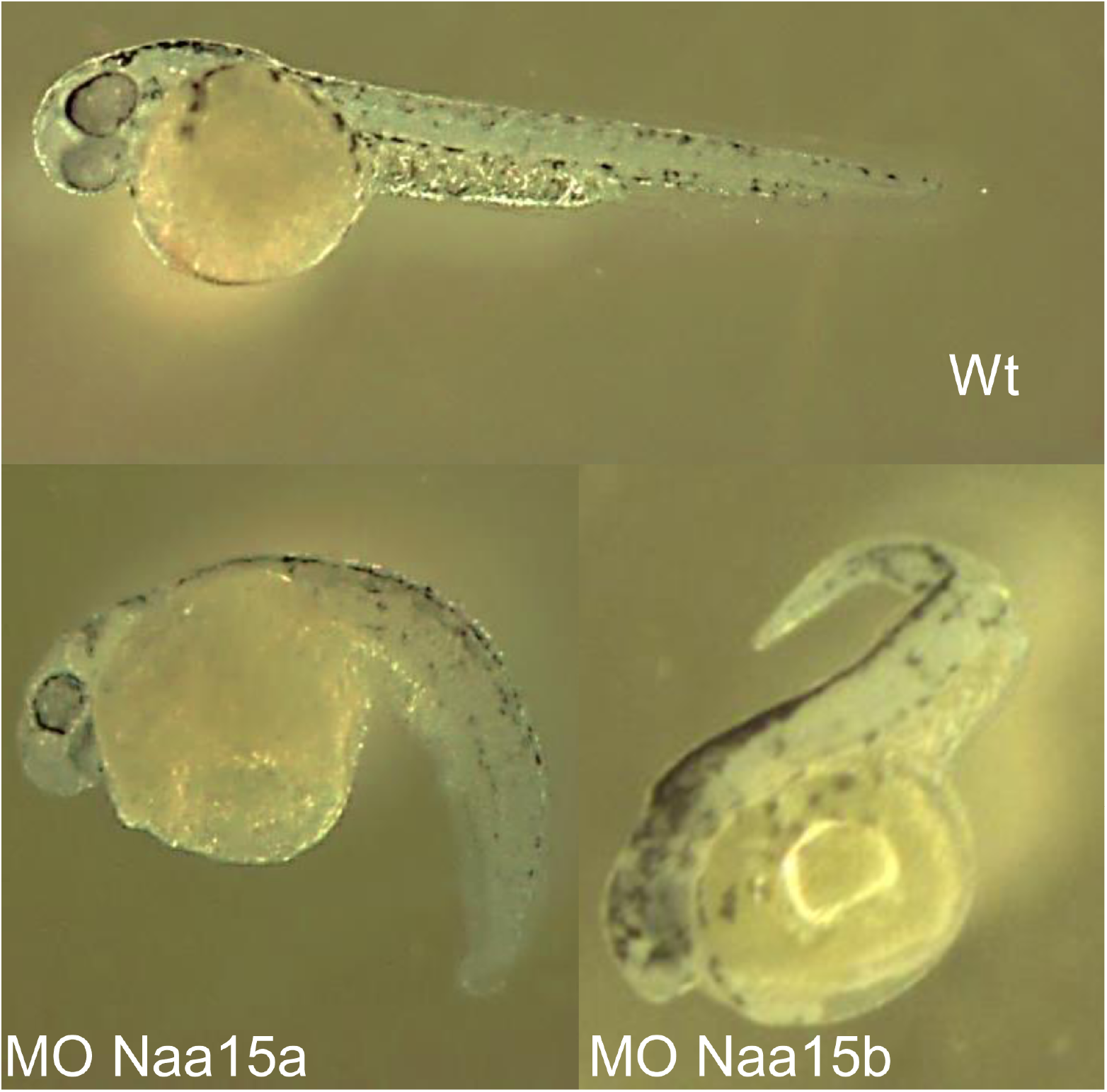
*Naa15* knockdown induce curvature of the body. Naa15 KD induces curvature of the body. Most of the eggs injected with anti-*naa15* morpholino gave rise to embryos with curvature of the trunk and reduction of body size. Injection of mismatch morpholino do not induce apparent phenotype. The majority of the *naa15a* morphants presents reduction of body size and moderate curvature of the trunk (bottom left) whereas most of the *naa15b* morphant presents greater curvature (bottom right).

Co-injection with anti-*naa15a* and anti-*naa15b* morpholino was also performed. Most of the embryos co-injected with the two morpholino died in the first 24 hours after fertilization. The survivors *naa15a* + *naa15b* morphants presented similar phenotypes to embryos injected with anti-*naa15a* or anti-*naa15b* alone (C1 and C2 phenotypes).

To confirm that the observed phenotypes did not result from unspecific induction of apoptosis (31), the experiments were replicated with embryos co-injected with anti-*naa15a* and anti-p53 morpholinos. In all experiments, the *naa15a/p53* morphants present the same phenotype than the embryos injected with anti-*naa15a* or anti-*naa15b* morpholinos alone.

### naa15a and naa15b knockdown in zebrafish led to intersegmental boundary defect

Confocal microscopy of the morphants after staining of the nuclei and the plasma membrane and connective tissue revealed defect in the segmentation of the C1 and C2 embryos as compare to control mismatch *naa15* morphants. Wheat Germ Agglutinin (WGA) conjugate with alexa488 was used for the plasma membrane and connective tissue staining. This protein is a lectin that selectively binds to N-acetylglucosamine and N-acetylneuraminic acid (sialic acid) residues and allows fast, and convenient methodology for connective tissue, and plasma membrane visualization (32). This methodology allowed us to shown that C1 and C2 embryos did not exhibit the usual chevron shape segmentation of the myotomes (Fig 5) (33). Further, the myotome of the *naa15a* and *naa15b* morphants presented a lack or reduction of the horizontal myoseptum and interruption of myotome boundaries along the dorso/ventral and medio/lateral axis (Fig 5). Those interruptions were observed in all the tested C1 and C2 embryos (n>15) in about 30% of the myotome boundaries. They formed holes through which some of the myofibres, displaying twice the normal fibre length, stretched out (figure 5, figure 6). Some of those longer myofibres could contain up to 11 nuclei but altogether, their mean number of nuclei per fibres was not significantly increased as compared to normal size fibres (not shown).

**Fig5.**
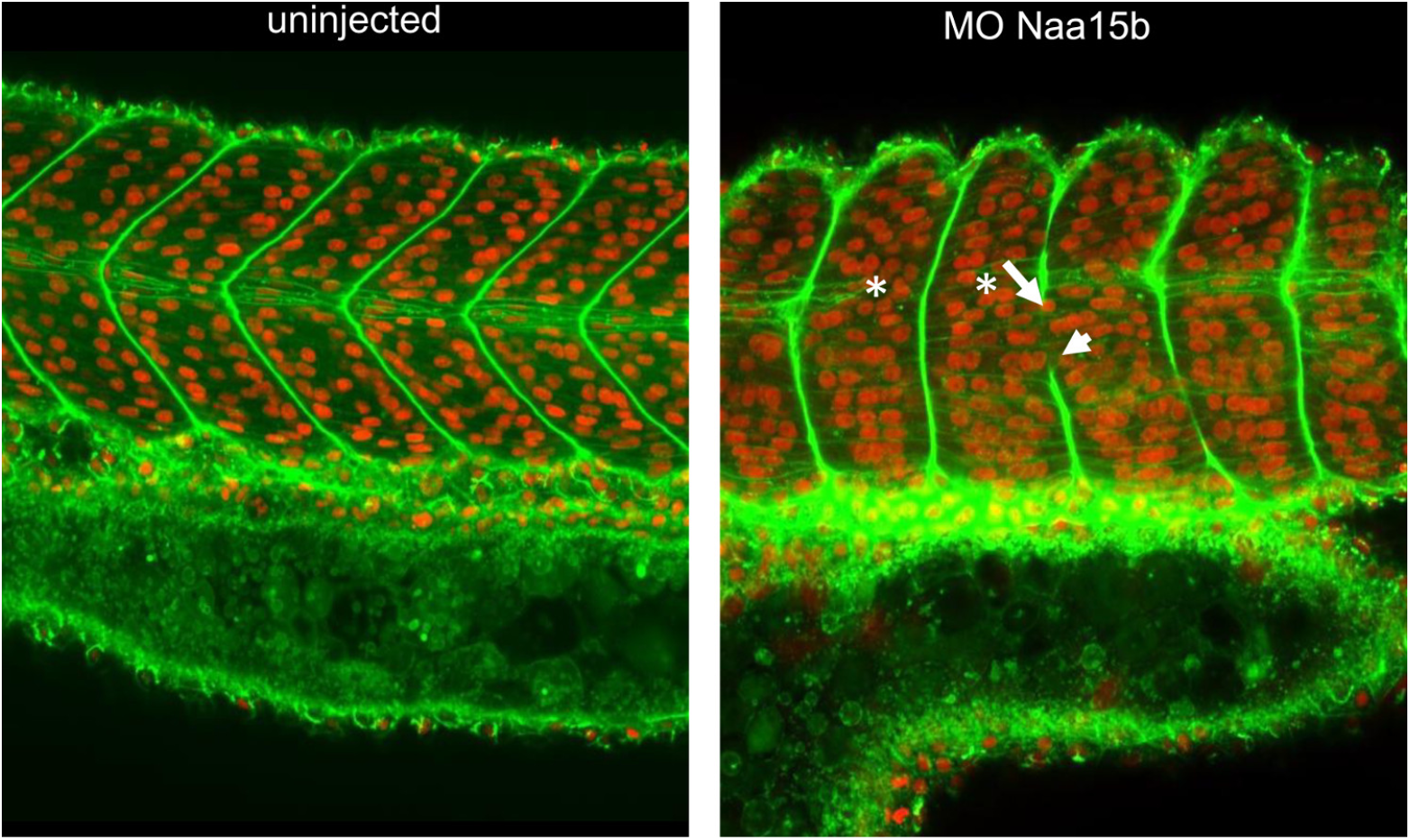
*Naa15* knockdown induce myotome defect. *naa15* KD induces myotomal boundary defects. The embryos were stained with TOPRO 3 (nuclei in red) and WGA-alexa488 (plasma membranes and myotome boundaries in green). Confocal microscopy of the group 1 and 2 embryos showed a loss of the classical chevron shaped segments with a lack or reduction of the horizontal myoseptum (asterisk) and interruption of myotome boundaries along the dorso/ventral and medio/lateral axis (arrow). Few myofibres stretch out within two somites/myotomes dues to interruption in somite boundaries (arrow head).

**Fig6.**
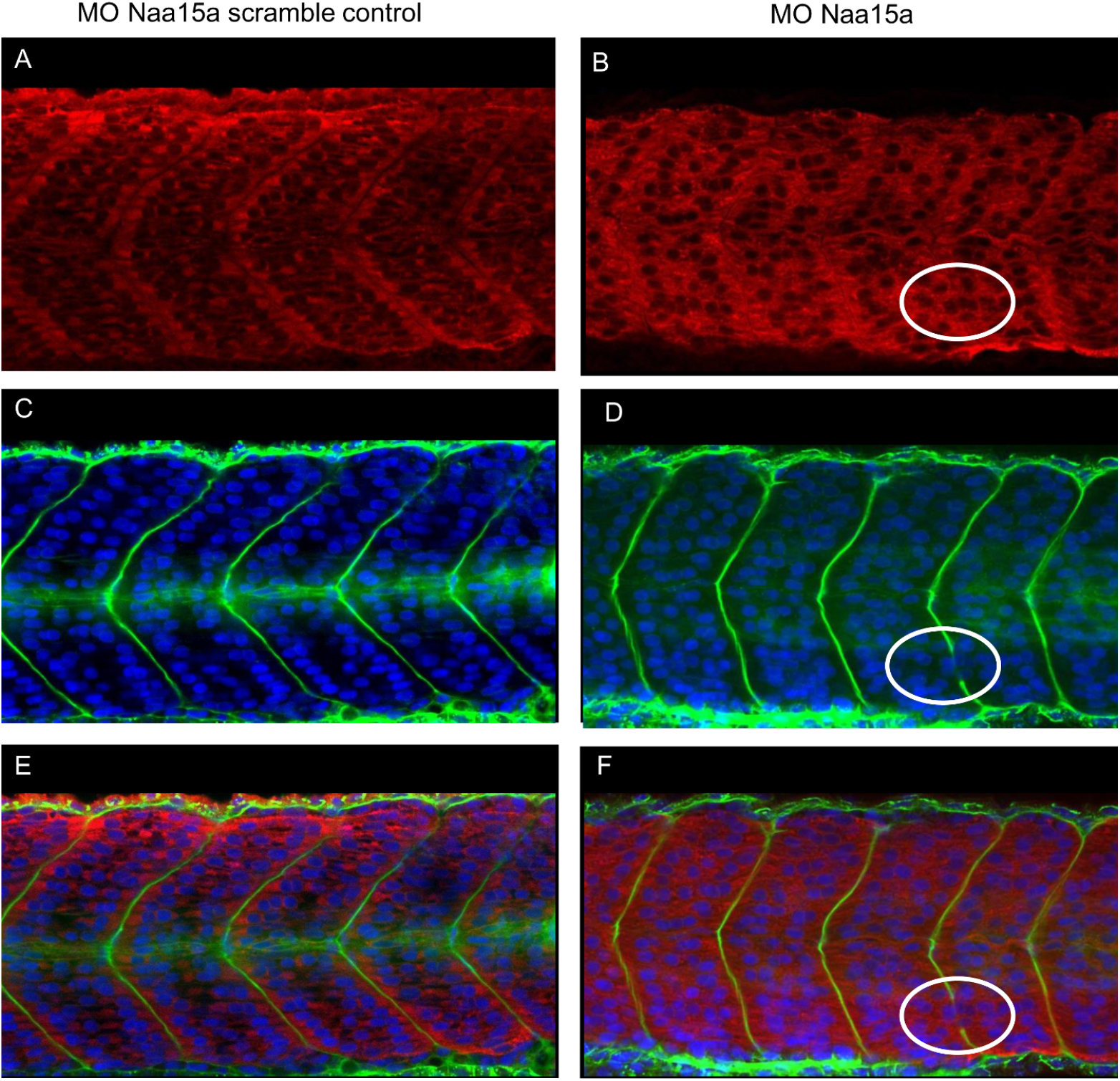
*Naa15* knockdown induce defect in the myosine organization. *naa15 KD induce defect in myosin organisation*. Confocal microscopy of morphants after staining with TOPRO 3 (nuclei in blue), WGA-alexa488 (plasma membranes and myotome boundaries in green) (panel A and B), and immunocytofluoresence staining with an antibody against myosin heavy chain (MF20, myosin in red) (panel C and D), revealed a disorganization of the myosin localization in the naa15 morphants (D) as compared to embryos injected with *naa15* mismatch *naa15a* morpholino (C). Some long myofibres project beyond the intersegmental boundaries (white circle).

### naa15a and naa15b knockdown in zebrafish led to a reorganization of the myosin pattern

In the C1 an C2 *naa15a* and *naa15b* morphants, the myosin proteins had not the same expression pattern than in the wild type embryos, as shown by myosins immunostaining performed using MF20 antibody. Myosin is mainly located at the peripheries of the wildtype control embryos myofibres, close to the myotome boundary (Fig 6A, E) whereas in the C1 and C2 morphants, it was uniformly present within the whole fibre (Fig 6B, F). This abnormal myosin localization was also observed in long fibre spanning the intersegmental boundaries (Fig 6B, F).

## 4. Discussion

The fusion of muscle progenitor cells is one of the critical steps in muscle formation and regeneration. This process requires several steps including cell recognition, adhesion, and membrane fusion. To identify new genes implicated in myocyte fusion in vertebrates, we undertook an siRNA screen in c2c12 myoblasts followed by *in vivo* functional studies of relevant candidate genes in zebrafish.

Among the tested genes, we identified *Naa15* as an inhibitor of c2c12 fusion since its knockdown enhanced fusion of myoblasts *in vitro. Naa15* is part of the family of N-terminal acetyltransferase subunits. The NAA15 protein is highly expressed during embryogenesis (34–36) and it binds to the catalytic subunit Naa10 to form the NatA complex (for review of NAT complex see (36)). In mammals, the NatA complex interact with various substrates and is implicated in a broad range of cellular processes from cell growth to cellular differentiation (35–44). The major function of the NatA complex is the proteins N-terminal acetylation (36). The N-terminal acetylatation has various consequence for a protein: it could determine the subcellular localization (45–48), module the protein-protein interactions (49,50) and is also crucial for protein folding. Neither Naa15 or the NatA complex is currently described to have a function during myogenesis. Nevertheless, other N-terminal acetyltransferase was reported to play a key role in tropomyosin-actin complex formation, increasing actin binding, and promoting the regulation of specific myosin activity (51,52).

Our *in silico* analysis revealed that the *Naa15* gene is found in two copies (*naa15a* and *naa15b*) in the zebrafish genome as a result of the Teleost Genome Duplication (TGD) (53). In situ hybridization analysis indicated that *naa15a* and *naa15b* were expressed in somites at 24h post fecundation, when myoblast fusion occurred (16). To further assess the role of *Naa15* in myogenesis, we undertook knockdown experiments in zebrafish using morpholinos. We observed that zebrafish embryos injected with anti *naa15* Morpholinos didn’t exhibit classical chevron-shaped myotomes. Further, interruptions in the intersegmental boundaries allowing long myofibres to span over two (rarely 3) segments. This phenotype is reminiscent with those observed after the knockdown of genes encoding components of the Notch signalling pathway, especially *her1* and *her7* (54,55).

NAA15 is known to be a binding partner of cortactin [31], a protein regulating the F-actin polymerization and as such could be involved in process such as migration, permeability or elongation of cells. The knockdown of *Naa15* in retinal endothelial cells induces activation of the c-SRC kinase resulting in the phosphorylation and activation of the cortactin by a still unknown mechanism (39). This activation of cortactin resulting of the *naa15a* or *naa15b* knockdown could be partially responsible of the phenotype we observed. Indeed, increasing cell permeability, adhesion and migration could lead to the *in vitro* enhancement of myoblast fusion, and the presence of longer myofibres *in vivo*. Nevertheless, we did not detect any significant modifications in cells migration or adhesion after *Naa15* knockdown (not shown).

In conclusion our results showed that *Naa15* not only inhibits c2c12 myoblast fusion *in vitro*, but also is expressed in zebrafish developing myotome where it appears to be essential for proper myotome formation. Further research is needed to decipher the possible functional link between *Naa15* activity and the notch pathway or the cortactin activity that could explain the phenotype of zebrafish embryos injected with anti *naa15* morpholinos. Altogether the better understanding of the acetylation process leading to the formation and reparation of muscle fibres will be useful to enhance muscle repair therapy.

## Acknowledgments

This project and O. Monestier fellowship were supported by the French National Research Agency (ANR-12-JSV7-0001-01). The fellowship of Aurélie Landemaine was supported by INRA PHASE and the Région Bretagne. The funders had no role in study design, data collection and analysis, decision to publish, or preparation of the manuscript. We also thank A. Patinote for zebrafish husbandry and the supply of eggs for microinjection and R Le Guével for automatic acquisition of pictures on the ImPACcell platform (http://imagerie-puces-a-cellules.univ-rennes1.fr).

## References

1. Tajbakhsh S. Skeletal muscle stem cells in developmental versus regenerative myogenesis. J Intern Med. 2009 Oct;266(4):372-89.

2. Rochlin K, Yu S, Roy S, Baylies MK. Myoblast fusion: when it takes more to make one. Dev Biol. 2010 May 1;341(1):66-83.

3. Strünkelnberg M, Bonengel B, Moda LM, Hertenstein A, de Couet HG, Ramos RG, et al. rst and its paralogue kirre act redundantly during embryonic muscle development in Drosophila. Development. 2001 Nov;128(21):4229-39.

4. Ruiz-Gómez M, Coutts N, Price A, Taylor MV, Bate M. Drosophila dumbfounded: a myoblast attractant essential for fusion. Cell. 2000 Jul 21;102(2):189-98.

5. Bour BA, Chakravarti M, West JM, Abmayr SM. Drosophila SNS, a member of the immunoglobulin superfamily that is essential for myoblast fusion. Genes Dev. 2000 Jun 15;14(12):1498-511.

6. Dworak HA, Charles MA, Pellerano LB, Sink H. Characterization of Drosophila hibris, a gene related to human nephrin. Development. 2001 Nov;128(21):4265-76.

7. Artero RD, Castanon I, Baylies MK. The immunoglobulin-like protein Hibris functions as a dose-dependent regulator of myoblast fusion and is differentially controlled by Ras and Notch signaling. Development. 2001 Nov;128(21):4251-64.

8. Shelton C, Kocherlakota KS, Zhuang S, Abmayr SM. The immunoglobulin superfamily member Hbs functions redundantly with Sns in interactions between founder and fusion-competent myoblasts. Development. 2009 Apr;136(7):1159-68.

9. Srinivas BP, Woo J, Leong WY, Roy S. A conserved molecular pathway mediates myoblast fusion in insects and vertebrates. Nat Genet. 2007 Jun;39(6):781-6.

10. Sohn RL, Huang P, Kawahara G, Mitchell M, Guyon J, Kalluri R, et al. A role for nephrin, a renal protein, in vertebrate skeletal muscle cell fusion. Proc Natl Acad Sci USA. 2009 Jun 9;106(23):9274-9.

11. Millay DP, O’Rourke JR, Sutherland LB, Bezprozvannaya S, Shelton JM, Bassel-Duby R, et al. Myomaker is a membrane activator of myoblast fusion and muscle formation. Nature. 2013 Jul 18;499(7458):301-5.

12. Landemaine A, Rescan P-Y, Gabillard J-C. Myomaker mediates fusion of fast myocytes in zebrafish embryos. Biochem Biophys Res Commun. 2014 Sep 5;451(4):480-4.

13. Quinn ME, Goh Q, Kurosaka M, Gamage DG, Petrany MJ, Prasad V, et al. Myomerger induces fusion of non-fusogenic cells and is required for skeletal muscle development. Nat Commun. 2017 Jun 1;8:15665.

14. Bi P, Ramirez-Martinez A, Li H, Cannavino J, McAnally JR, Shelton JM, et al. Control of muscle formation by the fusogenic micropeptide myomixer. Science. 2017 21;356(6335):323-7.

15. Zhang W, Roy S. Myomaker is required for the fusion of fast-twitch myocytes in the zebrafish embryo. Dev Biol. 2017 01;423(1):24-33.

16. Powell GT, Wright GJ. Jamb and jamc are essential for vertebrate myocyte fusion. PLoS Biol. 2011 Dec;9(12):e1001216.

17. Schwander M, Leu M, Stumm M, Dorchies OM, Ruegg UT, Schittny J, et al. Beta1 integrins regulate myoblast fusion and sarcomere assembly. Dev Cell. 2003 May;4(5):673-85.

18. Massarwa R’ada, Carmon S, Shilo B-Z, Schejter ED. WIP/WASp-based actin-polymerization machinery is essential for myoblast fusion in Drosophila. Dev Cell. 2007 Apr;12(4):557-69.

19. Richardson BE, Beckett K, Nowak SJ, Baylies MK. SCAR/WAVE and Arp2/3 are crucial for cytoskeletal remodeling at the site of myoblast fusion. Development. 2007 Dec;134(24):4357-67.

20. Schäfer G, Weber S, Holz A, Bogdan S, Schumacher S, Müller A, et al. The Wiskott-Aldrich syndrome protein (WASP) is essential for myoblast fusion in Drosophila. Dev Biol. 2007 Apr 15;304(2):664-74.

21. Kim S, Shilagardi K, Zhang S, Hong SN, Sens KL, Bo J, et al. A critical function for the actin cytoskeleton in targeted exocytosis of prefusion vesicles during myoblast fusion. Dev Cell. 2007 Apr;12(4):571-86.

22. Gildor B, Massarwa R’ada, Shilo B-Z, Schejter ED. The SCAR and WASp nucleation-promoting factors act sequentially to mediate Drosophila myoblast fusion. EMBO Rep. 2009 Sep;10(9):1043-50.

23. Moore CA, Parkin CA, Bidet Y, Ingham PW. A role for the Myoblast city homologues Dock1 and Dock5 and the adaptor proteins Crk and Crk-like in zebrafish myoblast fusion. Development. 2007 Sep;134(17):3145-53.

24. Laurin M, Fradet N, Blangy A, Hall A, Vuori K, Côté J-F. The atypical Rac activator Dockl80 (Dock1) regulates myoblast fusion in vivo. Proc Natl Acad Sci USA. 2008 Oct 7;105(40):15446-51.

25. Vasyutina E, Martarelli B, Brakebusch C, Wende H, Birchmeier C. The small G-proteins Rac1 and Cdc42 are essential for myoblast fusion in the mouse. Proc Natl Acad Sci USA. 2009 Jun 2;106(22):8935-40.

26. Gruenbaum-Cohen Y, Harel I, Umansky K-B, Tzahor E, Snapper SB, Shilo B-Z, et al. The actin regulator N-WASp is required for muscle-cell fusion in mice. Proc Natl Acad Sci USA. 2012 Jul 10;109(28):11211-6.

27. Hochreiter-Hufford AE, Lee CS, Kinchen JM, Sokolowski JD, Arandjelovic S, Call JA, et al. Phosphatidylserine receptor BAI1 and apoptotic cells as new promoters of myoblast fusion. Nature. 2013 May 9;497(7448):263-7.

28. Park D, Tosello-Trampont A-C, Elliott MR, Lu M, Haney LB, Ma Z, et al. BAI1 is an engulfment receptor for apoptotic cells upstream of the ELM0/Dock180/Rac module. Nature. 2007 Nov 15;450(7168):430-4.

29. Hamoud N, Tran V, Croteau L-P, Kania A, Côté J-F. G-protein coupled receptor BAI3 promotes myoblast fusion in vertebrates. Proc Natl Acad Sci USA. 2014 Mar 11;111(10):3745-50.

30. Thisse C, Thisse B. High-resolution in situ hybridization to whole-mount zebrafish embryos. Nat Protoc. 2008;3(1):59-69.

31. Bedell VM, Westcot SE, Ekker SC. Lessons from morpholino-based screening in zebrafish. Brief Funct Genomics. 2011 Jul;10(4):181-8.

32. Kostrominova TY Application of WGA lectin staining for visualization of the connective tissue in skeletal muscle, bone, and ligament/tendon studies. Microsc Res Tech. 2011 Jan;74(1):18-22.

33. Rost F, Eugster C, Schröter C, Oates AC, Brusch L. Chevron formation of the zebrafish muscle segments. J Exp Biol. 2014 Nov 1;217(Pt 21):3870-82.

34. Gendron RL, Adams LC, Paradis H. Tubedown-1, a novel acetyltransferase associated with blood vessel development. Dev Dyn. 2000 Jun;218(2):300-15.

35. Sugiura N, Adams SM, Corriveau RA. An evolutionarily conserved N-terminal acetyltransferase complex associated with neuronal development. J Biol Chem. 2003 Oct 10;278(41):40113-20.

36. Kalvik TV, Arnesen T. Protein N-terminal acetyltransferases in cancer. Oncogene. 2013 Jan 17;32(3):269-76.

37. Gendron RL, Good WV, Adams LC, Paradis H. Suppressed expression of tubedown-1 in retinal neovascularization of proliferative diabetic retinopathy. Invest Ophthalmol Vis Sci. 2001 Nov;42(12):3000-7.

38. Paradis H, Liu C-Y, Saika S, Azhar M, Doetschman T, Good WV, et al. Tubedown-1 in remodeling of the developing vitreal vasculature in vivo and regulation of capillary outgrowth in vitro. Dev Biol. 2002 Sep 1;249(1):140-55.

39. Paradis H, Islam T, Tucker S, Tao L, Koubi S, Gendron RL. Tubedown associates with cortactin and controls permeability of retinal endothelial cells to albumin. J Cell Sci. 2008 Jun 15;121(Pt 12):1965-72.

40. Geissenhöner A, Weise C, Ehrenhofer-Murray AE. Dependence of ORC silencing function on NatA-mediated Nalpha acetylation in Saccharomyces cerevisiae. Mol Cell Biol. 2004 Dec;24(23):10300-12.

41. Wall DS, Gendron RL, Good WV, Miskiewicz E, Woodland M, Leblanc K, et al. Conditional knockdown of tubedown-1 in endothelial cells leads to neovascular retinopathy. Invest Ophthalmol Vis Sci. 2004 Oct;45(10):3704-12.

42. Wang X, Connelly JJ, Wang C-L, Sternglanz R. Importance of the Sir3 N terminus and its acetylation for yeast transcriptional silencing. Genetics. 2004 Sep;168(1):547-51.

43. Asaumi M, Iijima K, Sumioka A, Iijima-Ando K, Kirino Y, Nakaya T, et al. Interaction of N-terminal acetyltransferase with the cytoplasmic domain of beta-amyloid precursor protein and its effect on A beta secretion. J Biochem. 2005 Feb;137(2):147-55.

44. Myklebust LM, Van Damme P, Støve SI, Dörfel MJ, Abboud A, Kalvik TV, et al. Biochemical and cellular analysis of Ogden syndrome reveals downstream Nt-acetylation defects. Hum Mol Genet. 2015 Apr 1;24(7):1956-76.

45. Behnia R, Panic B, Whyte JRC, Munro S. Targeting of the Arf-like GTPase Arl3p to the Golgi requires N-terminal acetylation and the membrane protein Sys1p. Nat Cell Biol. 2004 May;6(5):405-13.

46. Setty SRG, Strochlic TI, Tong AHY, Boone C, Burd CG. Golgi targeting of ARF-like GTPase Arl3p requires its Nalpha-acetylation and the integral membrane protein Sys1p. Nat Cell Biol. 2004 May;6(5):414-9.

47. Behnia R, Barr FA, Flanagan JJ, Barlowe C, Munro S. The yeast orthologue of GRASP65 forms a complex with a coiled-coil protein that contributes to ER to Golgi traffic. J Cell Biol. 2007 Jan 29;176(3):255-61.

48. Forte GMA, Pool MR, Stirling CJ. N-terminal acetylation inhibits protein targeting to the endoplasmic reticulum. PLoS Biol. 2011 May;9(5):e1001073.

49. Scott DC, Monda JK, Bennett EJ, Harper JW, Schulman BA. N-terminal acetylation acts as an avidity enhancer within an interconnected multiprotein complex. Science. 2011 Nov 4;334(6056):674-8.

50. Monda JK, Scott DC, Miller DJ, Lydeard J, King D, Harper JW, et al. Structural conservation of distinctive N-terminal acetylation-dependent interactions across a family of mammalian NEDD8 ligation enzymes. Structure. 2013 Jan 8;21(1):42-53.

51. Polevoda B, Cardillo TS, Doyle TC, Bedi GS, Sherman F. Nat3p and Mdm20p are required for function of yeast NatB Nalpha-terminal acetyltransferase and of actin and tropomyosin. J Biol Chem. 2003 Aug 15;278(33):30686-97.

52. Coulton AT, East DA, Galinska-Rakoczy A, Lehman W, Mulvihill DP. The recruitment of acetylated and unacetylated tropomyosin to distinct actin polymers permits the discrete regulation of specific myosins in fission yeast. J Cell Sci. 2010 Oct 1;123(Pt 19):3235-43.

53. Jaillon O, Aury J-M, Wincker P. “Changing by doubling”, the impact of Whole Genome Duplications in the evolution of eukaryotes. C R Biol. 2009 Mar;332(2-3):241-53.

54. Henry CA, Urban MK, Dill KK, Merlie JP, Page MF, Kimmel CB, et al. Two linked hairy/Enhancer of split-related zebrafish genes, her1 and her7, function together to refine alternating somite boundaries. Development. 2002 Aug;129(15):3693-704.

55. Henry CA, McNulty IM, Durst WA, Munchel SE, Amacher SL. Interactions between muscle fibers and segment boundaries in zebrafish. Dev Biol. 2005 Nov 15;287(2):346-60.

